# Label-free real-time imaging of mitochondrial matrix volume changes and permeability transition in living cells

**DOI:** 10.64898/2026.05.15.725497

**Authors:** Yaw Akosah, Ioannis Azoidis, Dane Jensen, Paolo Bernardi, Evgeny Pavlov

## Abstract

Along with the membrane potential and respiration, mitochondrial matrix volume is a critical parameter that determines mitochondrial function. Mitochondria undergo constant changes in matrix volume and cristae dynamics, and in processes that are critical for normal metabolic rates and pathophysiological responses. Changes in matrix volume cannot be easily measured by conventional fluorescence imaging techniques due to the size of the sub-organellar structures, which are below resolution. This challenge was successfully resolved in studies of isolated mitochondria with the use of scattered light. Here we use dark-field imaging, which relies on scattered light contrast, to measure matrix volume dynamics in living cells. We demonstrate that mitochondrial volume changes can be easily detected as changes in intensity of the scattered light following matrix volume modulation with K^+^ ionophores or by onset of the permeability transition. Specifically, we found that stimulation of K^+^ influx leads to increase of mitochondrial matrix volume while stimulation of K^+^ efflux leads to matrix shrinkage, and that activation of the permeability transition leads to high-amplitude mitochondrial swelling in wild-type but not in cells lacking subunit c of ATP synthase. These results directly demonstrate the dynamic nature of mitochondrial matrix volume and its link to physiological and pathological ion transport.

## Main

Mitochondria are intracellular organelles bound by inner and outer membranes. The inner mitochondrial membrane forms multiple folds, generating extensive cristae that expand the surface area. This dense packaging results in the ability of mitochondria to scatter light, a property that has been widely used to study mitochondrial volume changes and inner membrane dynamics ^1^. In a typical experiment, a suspension of isolated mitochondria is exposed to light and the scattered component is measured. The intensity of scattered light changes with the change in cristae morphology and matrix light scattering properties, with an increase in mitochondrial matrix volume leading to a decrease in light scattering. This approach has been used successfully to measure mitochondrial K^+^ transport, where K^+^ influx causes osmotic matrix swelling ^2^; and opening of the mitochondrial permeability transition pore (PTP), which causes high amplitude swelling and a dramatic drop in the intensity of scattered light ^3^. This sensitive method has so far been applied only to populations of isolated mitochondria. To expand the method to living cells, we applied a dark-field imaging technique where non-scattering objects (like the cytoplasm) appear as dark regions while scattering objects (like mitochondria) appear as bright bodies ^4^. By manipulating mitochondrial volume with well-characterized K^+^ ionophores we demonstrate that this method allows label–free identification of mitochondria and monitoring of their matrix volume changes. Further, we applied this method to monitor the consequences of PTP opening in wild-type cells and in cells where the c subunit of mitochondrial ATP synthase had been ablated. Our data highlight the dynamic nature of mitochondrial matrix volume and demonstrate that dark-field imaging is a powerful tool to study mitochondrial ion transport and volume changes in situ.

## Results

### Detection of mitochondria in living cells using dark-field imaging

The principle of the dark-field imaging technique is very similar to that of light scattering in isolated mitochondrial populations. In the case of isolated mitochondria the detector captures the light scattered by the mitochondrial suspension and the intensity of the scattered light changes depending on the mitochondrial volume changes, with decreased scattering as matrix volume increases (Fig. 1, left panel). In the case of dark-field imaging the opaque ring positioned in front of the light source (Fig. 1, right panel) is adjusted in such a way that only the light scattered by the sample is collected, while transmitted light is not detected ^5^ (Fig. 1, right panel).

**Figure 1.**
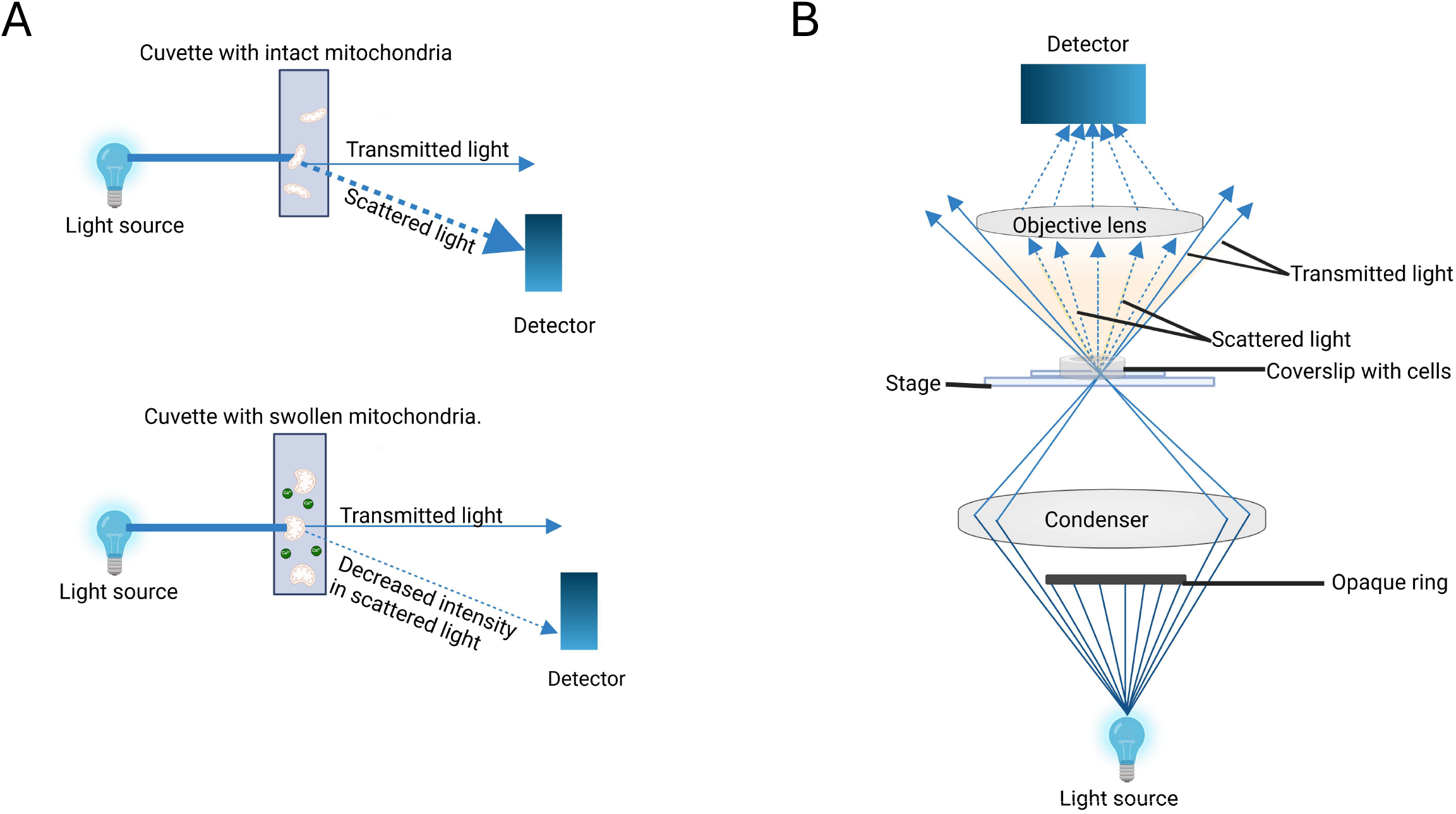
Schematic representation of the principles of sample detection in light-scattering and dark-field approaches. **A.**In light-scattering assay mitochondrial sample is incubated in recording solution and matrix volume changes measured as changes in the intensity of scattered light. **B**. In dark-field imaging an opaque ring is positioned in front of the light source to eliminate all transmitted light from the objective lens, thus allowing selective detection of light that was scattered by the biological sample.

First, we aimed to investigate how mitochondria can be viewed in the dark-field mode in living cultured cells. The dark field images of HEK and MEF cells indicate the presence of light-scattering regions (Fig. 2A,D). These regions include mitochondria, identified by staining with the membrane potential fluorescent probe TMRM (Fig. 2B,E), which colocalized with most of the cytoplasmic structures seen with the dark-field mode (Fig. 2C,F). Light scattering, but not TMRM staining, was also observed for nuclear structures unrelated to mitochondria. These results indicate that in our experimental setup mitochondria can be readily visualized using dark-field imaging. Importantly, cells contained regions where most of the light was scattered from mitochondria (Fig. 2G,H), and these were used for quantitative real-time studies of mitochondrial light-scattering properties as further detailed below.

**Figure 2.**
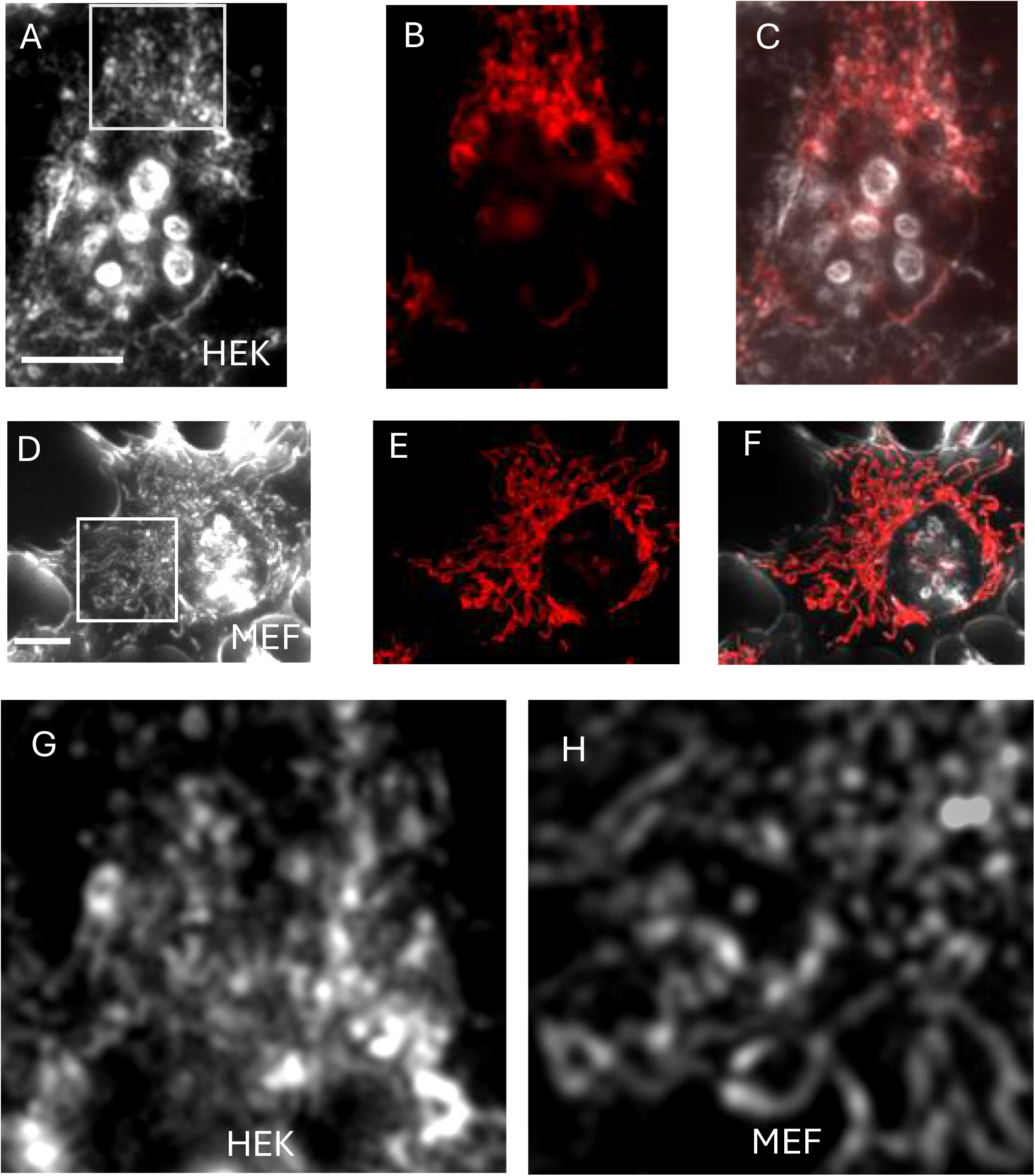
Visualization of mitochondria in living cells using dark-field imaging. **A.**Dark-field image of HEK cells; **B**. Same field as in (B) showing the TMRM fluorescent signal; **C**. Overlay of images A and B; **D**. Dark-field image of MEF cells; **E**. Same field as in (D) showing the TMRM fluorescent signal; **F**. Overlay of images D and E. **G**,**H**. Enlarged images of the areas boxed in panels A and D, respectively. Scale bars 10 µm.

### Real-time monitoring of mitochondrial volume changes induced by K^+^ fluxes

Experiments with isolated mitochondria indicate that their ability to scatter light decreases with the increase in mitochondrial volume. This property has been used extensively to study mitochondrial K^+^ transport. In experiments with suspensions of mitochondria, K^+^ uptake causes matrix osmotic swelling, which can be quantified in real time using light-scattering intensity measurements ^1^. Here, we tested whether similar measurements can be performed in mitochondria of living cells. Light scattering was monitored in dark-field images (Fig. 3A) and mitochondria were identified by labeling with TMRM (Fig. 3B). Mitochondrial K^+^ uptake was then induced with the K^+^-selective ionophore valinomycin, which caused disappearance of mitochondria from the dark-field images (Fig. 2Ai-iii) followed by a drop in the TMRM signal (Fig. 3Bi-iii). The time course of the changes of light scattering (Fig. 3C, left panel) and fluorescence (Fig. 3D, left panel) indicates that onset of swelling precedes depolarization. To further define the bases for the volume changes, we measured light scattering when valinomycin was added in the presence of nigericin, a selective K^+^-H^+^ exchanger that allows extrusion of K^+^ ions from the mitochondrial matrix preventing K^+^ accumulation and generation of osmotic force, diverting respiration to the futile cycling of K^+^ and H^+^ across the inner membrane ^6^. As predicted, under these conditions mitochondria depolarized immediately after the addition of valinomycin and light scattering remained constant (Fig. 3C,D, center panels). Finally, addition of nigericin alone, which causes mitochondrial K^+^ efflux and matrix shrinkage, induced a marked increase of light scattering (Fig. 3C, right panel) while membrane potential remained stable (Fig. 3D, right panel). All the effects reported in panels C and D of Fig. 3 were statistically significant (Fig. 3E). Electron microscopy imaging confirmed that the light scattering changes, or lack thereof, were due to the changes of in situ mitochondrial volume and morphology. Indeed, mitochondria in untreated cells contained intact and easily identifiable cristae, treatment with valinomycin caused increased mitochondrial volume and decreased cristae density, and nigericin caused matrix contraction with marked increase of cristae density (Fig. 3F and Suppl. Fig. 1). Overall, these findings indicate that the technique is able to monitor changes of mitochondrial matrix volume in either direction.

**Figure 3.**
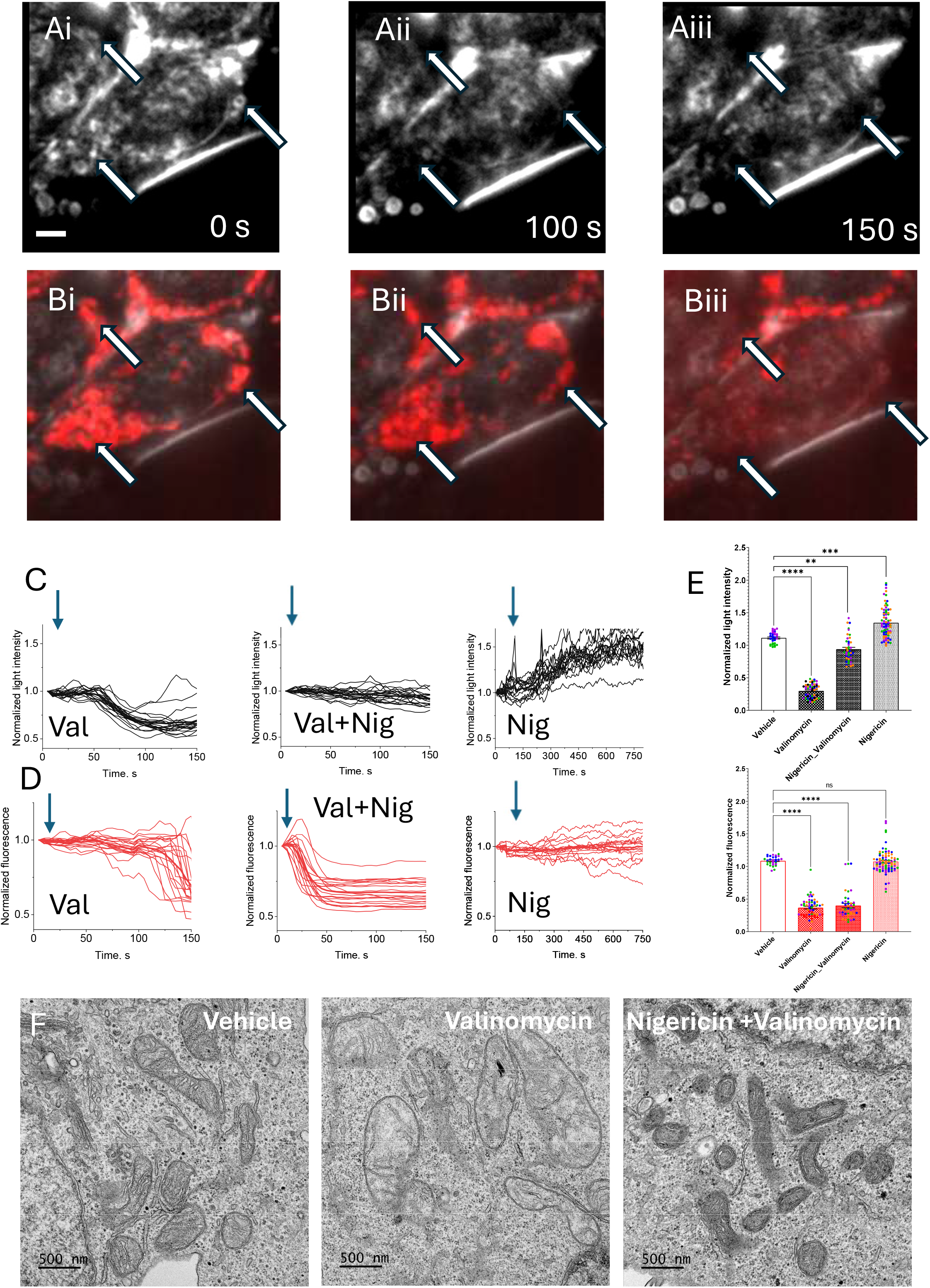
Change in mitochondrial light scattering in response to valinomycin, nigericin and their combination. **Ai-Aiii.=**Dark-field images of living HAP1 cells taken at the indicated time points following valinomycin addition. Mitochondrial regions are marked by arrows; **Bi-Biii**. The same images as in A were overlayed with TMRM signal, which was used to identify mitochondrial regions of the cell; scale bar 5 µm; **C**,**D**. Time course of scattered light (C) and TMRM intensity (D) following addition of valinomycin (left panels), valinomycin in the presence of nigericin (center panels) or nigericin alone (right panels). Time of addition shown by the blue arro; **E**. Relative drop in light scattering and TMRM intensities under various experimental conditions. The ratios were calculated by dividing intensities at the end and the beginning of experiments, same color points represent data from an individual culture slide (vehicle: N=3; n=30, valinomycin: N=5; n=56, vigericin+valinomycin: N=4 ; n=42, nigericin: N=4, n=81); **F**. Electron microscopy images of mitochondria following the addition of vehicle, valinomycin or valinomycin + nigericin. Note the mitochondrial swelling response to valinomycin and shrinkage response to nigericin.

### In situ detection of mitochondrial permeability transition pore opening

In isolated mitochondria the light-scattering assay has been used extensively to characterize Ca^2+^-induced opening of the PTP, a high-conductance non-selective pore. Long-lasting openings of the PTP disrupt mitochondrial function and eventually result in high-amplitude mitochondrial swelling, which can be detected as a decrease of light scattering. Here, we tested if PTP opening can be detected in the mitochondria of living cells using dark-field imaging. To induce PTP opening we used the Ca^2+^ ionophore ferutinin, which allows Ca^2+^ uptake and its accumulation inside the mitochondrial matrix. This accumulation results in membrane depolarization and permeabilization, which can be inhibited by cyclosporin A (CsA) ^7-9^, indicative of PTP opening. Within seconds of the addition of ferutinin, decreased light scattering (Fig. 4Ai-iv) and mitochondrial depolarization took place (Fig. 4Bi-iv), both events being consistent with PTP opening followed by high-amplitude swelling. The latter can be appreciated particularly well in the area expanded in Fig. 4Ci-iv and in the electron micrographs (Fig. 4D), and in the analysis of the time course of the detected changes in light scattering (Fig. 4E). In control experiments CsA addition alone did not cause mitochondrial depolarization or decreased light scattering (Suppl. Fig. 2). Notably, unlike the case of valinomycin addition when all mitochondria responded simultaneously, disappearance of the mitochondrial signal induced by ferutinin (which we interpret as due to PTP opening) occurred at different time points within different subpopulations of mitochondria (Fig. 4Ci-iv and Suppl. Fig. 3). This phenomenon is also evident from the highly heterogenous time course of light intensity changes measured in different mitochondrial regions (Fig. 4E, left panel). Remarkably, average of individual traces (Fig. 4E, green line) appears as a gradual loss of light scattering when the whole population of cells in taken into consideration. As a further test, we treated cells with the protonophore FCCP, which caused prompt depolarization but no changes in light scattering (Fig. 4F). Overall, the changes of light scattering and fluorescence intensity, or lack thereof, were statistically significant (Fig. 4G). Next, we utilized this novel approach to investigate the features of the permeability transition in cells lacking the c-subunit of the ATP synthase – a key component of the PTP. Previous experiments indicated that in the absence of the c-subunit Ca^2+^ accumulation still leads to opening of a CsA-sensitive channel in the inner mitochondrial membrane, and that this channel is significantly smaller than the PTP seen in wild-type cells ^10,11^. Consistent with these previous findings, we found that in cells lacking subunit c the addition of ferutinin led to membrane depolarization but not to mitochondrial swelling (Fig. 5). It should also be noted that in the case of mutant cells membrane depolarization occurred at a slower rate, compared to the wild-type; and that depolarization was inhibited by CsA, which is consistent with previous findings and with the idea that depolarization is due to opening of a low conductance or smaller pore. These results confirm that an assembled ATP synthase is required for induction of PTP opening and high-amplitude mitochondrial swelling.

**Figure 4.**
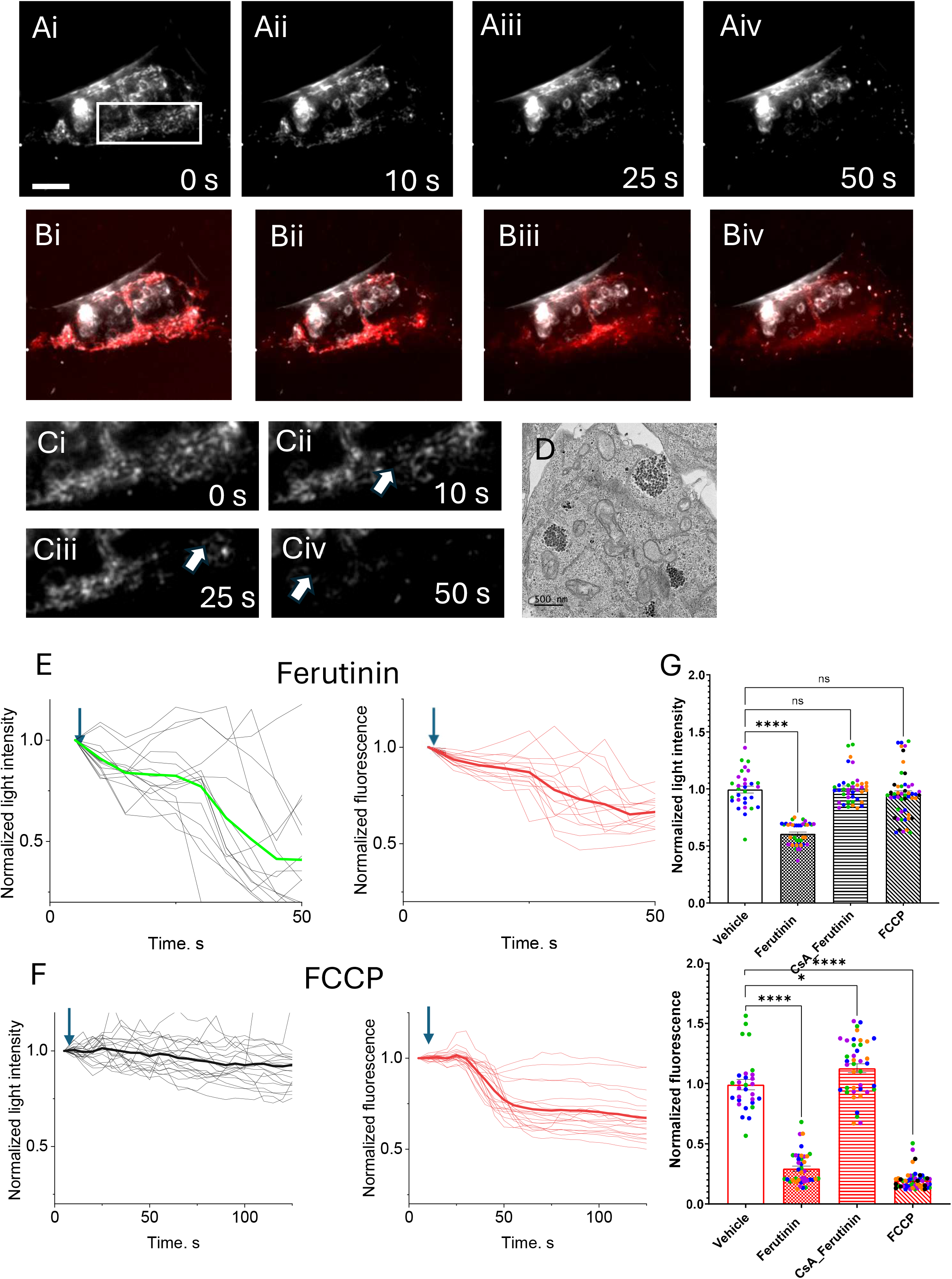
Mitochondrial swelling in intact cells following activation of the permeability transition pore. **A.**Dark-field images of living HAP1 cells taken at different time points following addition of ferutinin scale bar, 5 µm; **B**. Same images as in A overlayed with TMRM signal, which was used to identify mitochondrial regions of the cell; scale bar, 5 μM; **C**. Enlargement of the boxed area in panel A, note that mitochondrial signal drops sequentially at different regions (marked by arrows) at different timepoints; **D**. Electron microscopy images of mitochondria following ferutinin treatment; **E**,**F**. Time course of scattered light (left) and TMRM intensity (right) following addition of ferutinin (E) or FCCP (F). Time of addition is denoted by the blue arrow; **G**. Relative drop in light scattering and TMRM intensities following various additions. The ratios were calculated by dividing intensities at the end and the beginning of experiments, same color points represent data from an individual culture slide (Vehicle: N=3; n=30, Ferutinin: N=4 n=40, CsA + Ferutinin: N=4, n=40; FCCP: N=5, n=50).

**Figure 5.**
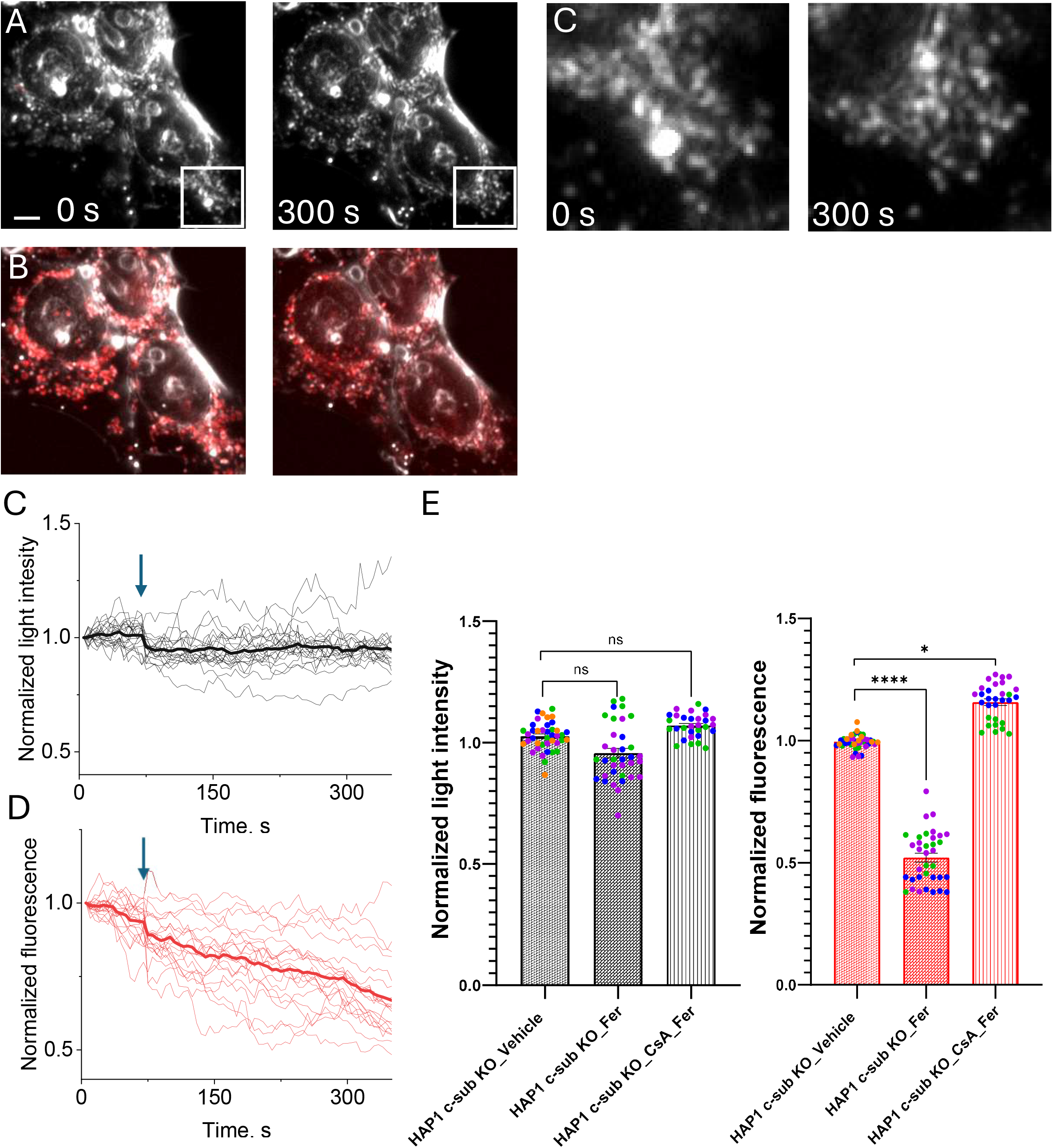
Lack of mitochondrial swelling in cells lacking subunit c of the ATP synthase. **A.**Dark-field images of living HAP1 c-subunit knock-out cells taken before and 300 s after ferutinin addition;scale bar, 5 µm; **B**. Same images as in A overlayed with the TMRM signal; **C**,**D**. Time course of the scattered light (left) and TMRM (right) intensity following addition of ferutinin (C) or FCCP (D). Time of addition shown by the blue arrow; **E**. Relative drop in light scattering and TMRM intensities following various additions, same color points represent data from an individual culture slide (HAP1 WT_Fer: N= 4, n=40; HAP1 c-sub KO: N=4, n=40; HAP1 c-sub KO_Fer: N=3 ; n=35).

## Discussion

The results described in the present paper demonstrate that dark-field imaging allows to detect mitochondria and to dynamically measure changes of their matrix volume in living cells. This method complements the use of fluorescent probes to measure mitochondrial volume in the cell but has several advantages. Fluorescent probes only resolve overall mitochondrial shape and provide information on total mitochondrial volume, which includes both the matrix and the intermembrane space ^12^. Furthermore, fluorescent probes may not distribute evenly in submitochondrial compartments, and they may be the source of perturbations such as phototoxicity, generation of reactive oxygen species and inhibition of respiration ^13^. The latter events obviously do not occur with the use of light scattering, which has the additional advantage of measuring matrix rather than total volume changes.

Under energized, coupled conditions mitochondrial K^+^ uptake occurs on the K^+^ uniporter and is driven by the K^+^ electrochemical gradient, while K^+^ release occurs on the K^+^-H^+^ exchanger and is largely driven by the H^+^ gradient. Our results suggest that the state of the mitochondrial matrix in situ corresponds to the orthodox state of Hackenbrock, which is characterized by an expanded matrix with well-organized cristae and an extremely limited intermembrane space ^14-16^. Consistently, the organelles displayed increased light-scattering (i.e. they underwent contraction) after the addition of the K^+^-H^+^ ionophore nigericin, assuming a morphology that appears to be closer to the condensed conformation of Hackenbrock; while they became nearly invisible (i.e. they underwent swelling) after the addition of the K^+^ ionophore valinomycin. These experiments provide an effective calibration of mitochondrial volume in situ (from maximally condensed to maximally swollen), a calibration that can be used as a standard reference to quantitatively evaluate the initial matrix volume and its changes in the protocols under study. It should be stressed that the light scattering measurements described here do not perturb mitochondrial function, and thus allow to reliably detect the changes of matrix volume that determine the changes in cristae shape.

Our study suggests that the dark-field method can be used to detect mitochondrial swelling in the context of PTP studies in intact cells. Traditionally, a drop in light-scattering induced by Ca^2+^ in a population of isolated mitochondria is considered a direct indication of PTP opening followed by solute diffusion. To mimic these conditions in intact cells we used the Ca^2+^ ionophore ferutinin, which has been previously shown to mediate matrix Ca^2+^ accumulation followed by CsA-sensitive membrane depolarization^7-9,17,18^. The refractive index imaging technique used in the present work demonstrates that ferutinin does induce CsA-sensitive permeabilization of the mitochondrial inner membrane, as judged by equilibration of the refractive index between cytoplasm and matrix regions, suggesting the nonselective nature of the permeability increase and supporting the contention that ferutinin causes opening of the PTP. Furthermore, our data provide additional support to the idea that the c-subunit of ATP synthase may be required for occurrence of the high-conductance PTP. Previous studies demonstrated that when challenged with ferutinin, mitochondria in c-subunit knock-out cells still depolarize in a CsA-sensitive process and yet the PTP high-conductance channel was not present ^8,10^, which is consistent with the lack of mitochondrial swelling in our experiments.

The ability to directly measure mitochondrial volume changes in living cells has important implications. This method allows to test if effects seen in model experiments with isolated mitochondria can also be observed in intact cells, and how these properties might differ. Indeed, volume changes in isolated mitochondria rely on media that cannot reproduce the composition of the cytoplasm, in turn limiting the predictions that can be made to the conditions occurring in situ. In addition to ions (mainly K^+^, Na^+^ and Cl^-^) the cytoplasm contains proteins, small molecules and other organelles that contribute to the osmotic and oncotic forces, and to the tethers that regulate mitochondrial volume. Our experiments may have provided the first direct demonstration that mitochondrial volume dynamics in living cells is determined by controlled K^+^ fluxes, i.e. K^+^ uptake by a uniport mechanism and K^+^ release via Mitchell’s K^+^-H^+^ exchanger ^19^; and that mitochondrial volume increases as a consequence of lasting PTP openings. Interestingly, valinomycin caused a gradual volume increase in all mitochondria, while activation of the PTP triggered volume increase in discrete groups of mitochondria. Irrespective of whether the latter pattern is due to gradual diffusion of the ionophore or to the existence of mitochondrial subpopulations with different thresholds for PTP opening, this finding is an important proof of concept that the technique can monitor volume changes of spatially resolved organelles.

## Materials and methods

### Cell cultures

Human HAP1 wild-type (WT) cells and their mutated variants, deficient in ATP synthase c-subunit (c-sub KO), were purchased from Horizon Discovery Biosciences Limited (Cambridge, UK) and maintained in accordance with the supplier’s protocol. Briefly, the cells were cultured in Iscove’s Modified Dulbecco’s Medium (IMDM), supplemented 1% Antibiotic Antimycotic Solution (Penicillin/Streptomycin/Amphotericin B; Sigma Aldrich), 10% heat-inactivated fetal bovine serum (GibcoTM), and 2 mM L-Glutamine (Agilent, USA). Cells were cultured in a humidified cell culture incubator under a 5% CO_2_ atmosphere at 37°C. Plating on coverslips and standard passaging were performed after the incubated cells reached a confluency of 70-90%. Cells were seeded onto poly-D-lysine–treated glass coverslips approximately 48 hours prior to imaging, allowing them to reach 70–90% confluency.

### Ionophores

Analytical grade ionophores and compounds used in this study were purchased from Sigma-Aldrich. Stock solutions of valinomycin (10 mM), nigericin (10 mM), ferutinin (10 mM), and carbonyl cyanide-*p*-trifluoromethoxyphenylhydrazone (FCCP; 10 mM) were prepared in DMSO, aliquoted, and stored at −20 °C. Working concentrations for all compounds were established through preliminary titration experiments in which incrementally increasing doses were applied until a reproducible response was observed. The mechanisms of action and final concentrations of ionophores used to treat cells are presented in Table 1.

| Ionophore | Concentration, μM | Mechanism of action |
| --- | --- | --- |
| Valinomycin | 1.0 | $K^+$ ionophore; selectively transports $K^+$ across the inner mitochondrial membrane, dissipating the $K^+$ gradient and promoting mitochondrial swelling<sup>20</sup>. |
| Nigericin | 5.0 | K<sup>+</sup>/H<sup>+</sup> antiporter; exchanges K<sup>+</sup> for H<sup>+</sup>, collapsing ΔpH and catalyzing K<sup>+</sup> efflux leading to mitochondrial contraction <sup>21,22</sup>. |
| Ferutinin | 20.0 | Ca<sup>2+</sup> ionophore; transports Ca<sup>2+</sup> into mitochondria, triggering Ca<sup>2+</sup> overload and inducing mPTP opening <sup>23</sup>. |
| FCCP | 1.0 | Protonophore (uncoupler); collapses mitochondrial membrane potential by carrying H<sup>+</sup> across the inner membrane depolarizing mitochondria <sup>24</sup>. |

### Mitochondrial Membrane Potential

Immediately before imaging, working dilutions were prepared in pre-warmed recording buffer consisting of HBSS supplemented with 40 nM tetramethylrhodamine methyl ester (TMRM), a cationic fluorophore used to monitor mitochondrial membrane potential (ΔΨm). Coverslips containing plated cells were transferred to an imaging chamber and gently rinsed twice with Hank’s Balanced Salt Solution (HBSS; Gibco) to remove residual culture medium. For dye loading, cells were incubated in recording buffer for 20 minutes at room temperature in the dark. After loading, cells were imaged, and real-time changes in ΔΨm were monitored continuously during ionophore treatment. TMRM was maintained in the recording buffer during all imaging to allow steady-state monitoring of mitochondrial membrane potential (ΔΨm).

### Swelling Assay

All mitochondrial swelling assays were conducted using HAP1 WT cells. Swelling was elicited using the K^+^ ionophore valinomycin and the electroneutral K^+^/H^+^ exchanger nigericin under three experimental conditions. In the first condition, cells were imaged under baseline recording for 50–75 sec, after which 1 μM valinomycin was applied to assess the effect of electrogenic K^+^ influx on mitochondrial light-scattering. In the second condition, cells were pre-incubated with 5 μM nigericin for 10 min to establish electroneutral K^+^/H^+^ exchange prior to imaging. Following baseline acquisition, 1 μM valinomycin was added to evaluate how pre-established K^+^/H^+^ exchange prevents valinomycin-induced mitochondrial swelling. In the final condition, 1 μM nigericin was added to initiate K^+^/H^+^ exchange across the inner mitochondrial membrane.

### PTP induction

To monitor PTP activation in HAP1 WT and HAP1 c-sub KO cells, we used ferutinin—a Ca^2+^-mobilizing ionophore frequently employed to trigger mitochondrial permeability transition. Ferutinin elevates cytosolic Ca^2+^ levels and drives its electrophoretic uptake into mitochondria, thereby creating conditions that favor PTP induction. In the first experimental condition, cells were imaged under baseline acquisition for 50–75 sec before the addition of 20 μM ferutinin to evaluate Ca^2+^-driven induction of PTP opening in both WT and mutant cell lines. In the second condition, cells were pre-treated with 2 μM CsA, a well-established inhibitor of PTP, for 5 min before imaging. After baseline recording, 20 μM ferutinin was introduced to determine how pharmacological inhibition of the pore alters Ca^2+^-induced mitochondrial swelling and depolarization.

### Dark-field and Fluorescent Imaging

To monitor mitochondrial light-scattering behavior in intact cells, we employed dark-field microscopy, an imaging approach in which the direct illumination path is excluded^5^. We used the Leica DMi8 Widefield fluorescent microscope equipped with a dark-field condenser. A Target Präparat of *Convallaria rhizome* (As3211), procured from Leica Microsystems, was used for aperture setup, as well as fluorescence and darkfield image calibration. Dark-field images (Light-scattering signals) together with TMRM fluorescence (TRITC Channel) were synchronously captured at 5-second intervals using a 40x objective. Following a 50–75-second baseline acquisition, ionophores were added at designated time points, and changes in light-scattering were monitored and quantified. Five micromolar FCCP was used to collapse the membrane potential, providing a reference point for normalization of the TMRM signal.

### Image processing

All microscopic images were exported as Multipage TIF files and analyzed in Fiji. ^25^To quantify mitochondria-specific light-scattering signals, dark-field images were spatially aligned and overlaid onto the corresponding TMRM fluorescence frames using the merge function in ImageJ. Only dark-field signals that exhibited clear co-localization with TMRM fluorescence were attributed to mitochondrial structures. Regions of interest (ROIs) corresponding to polarized, functionally intact mitochondria were manually selected based on the initial TMRM signal at t = 0 s. ROIs selection did not allow to track individual mitochondria as each ROI contained several organelles and corresponding fluorescent and scattered light intensity was average of the signal originating from the whole ROI. In our control experiments with vehicle (DMSO), fluorescence and light scattering didn’t change over the same time course, indicating that mitochondrial movement and bleaching didn’t contribute to the signal readout. For each time point, the mean intensity values of both the dark-field (scattered light) and TMRM signals within each ROI were extracted. Signal values were normalized to their respective baseline intensities at t = 0 s to allow comparison across cells and imaging sessions. A progressive decrease in normalized dark-field intensity within co-localized regions was interpreted as a reduction in mitochondrial light-scattering, consistent with matrix swelling and inner membrane permeabilization. A decline in the mitochondrial membrane potential was simultaneously evaluated by a decrease in TMRM fluorescence intensity.

### Transmission Electron Microscopy (TEM)

Prior to transmission electron microscopy (TEM) imaging, vehicle- and drug-treated cells were cultured in 35-mm glass-bottom dishes. Immediately following treatment, the recording buffer was removed, and cells were fixed with 1 mL of freshly prepared fixation solution containing 2% paraformaldehyde and 2.5% glutaraldehyde in 0.1 M sodium cacodylate buffer (pH 7.2–7.4). Dishes were gently rotated to ensure uniform exposure of all cells to the fixative and incubated at room temperature for 1 min. Fixed cells were harvested using a cell scraper, transferred into 1.5-mL microcentrifuge tubes, and centrifuged at 800 g for 30 s at room temperature. Tubes were subsequently rotated 180° and centrifuged for an additional 1 min to ensure complete pelleting of cells. Cell pellets were left undisturbed in fixation solution for 30 min prior to downstream processing. TEM imaging was performed at the NYU Langone Microscopy Laboratory using a JEOL JEM-1400 Flash Transmission Electron Microscope equipped with a Gatan Rio CMOS Camera.

TEM images were analyzed using ImageJ/Fiji software. Raw TEM micrographs (5000X) were imported as TIFF images and calibrated using the embedded scale bar prior to analysis. The scale was set by drawing a line across the scale bar using the straight-line tool, followed by selection of *Analyze* → *Set Scale* to convert pixel dimensions to micrometers (µm). Images were converted to 8-bit grayscale format, and brightness/contrast was adjusted uniformly across all experimental groups when necessary. Individual mitochondria were manually outlined using the freehand selection tool to ensure accurate segmentation of mitochondrial boundaries. Each traced mitochondrion was added to the Region of Interest (ROI) Manager for subsequent quantification. Morphometric measurements were obtained by selecting *Analyze* → *Set Measurements* and enabling the parameters for area and perimeter. Mitochondrial area (µm^2^) and perimeter (µm) were then quantified using the *Measure* function in ROI Manager. All measurements were performed using identical analysis settings across samples. For each experimental condition, multiple cells and mitochondria were analyzed from independent biological replicates. Quantitative data were exported as .csv files and analyzed statistically using GraphPad Prism (version 10.6.1).

### Statistical processing of data

GraphPad Prism software (version 10.6.1) was used for all statistical analyses. Data are presented as mean ± SEM. Statistical analyses were performed using a mixed-effects model with biological replicate treated as a random effect ^26^. Where appropriate, one-way ANOVA followed by Dunnet’s multiple comparisons test was applied to biological replicate-level summaries using. Statistical significance was defined as *p ≤ 0.05; **p ≤ 0.01; ***p ≤ 0.001; ****p ≤ 0.0001. The number of independent experiments (N) and the number of cells analyzed per experiment (n) are indicated in the corresponding sections of the manuscript and figure legends.

## Acknowledgement

We thank NYU Grossman School of Medicine DART Microscopy Laboratory for the consultation and assistance with EM work. This work was supported by NIH grant R35GM139615 (EP)

---

**Supplementary Figure 1.**
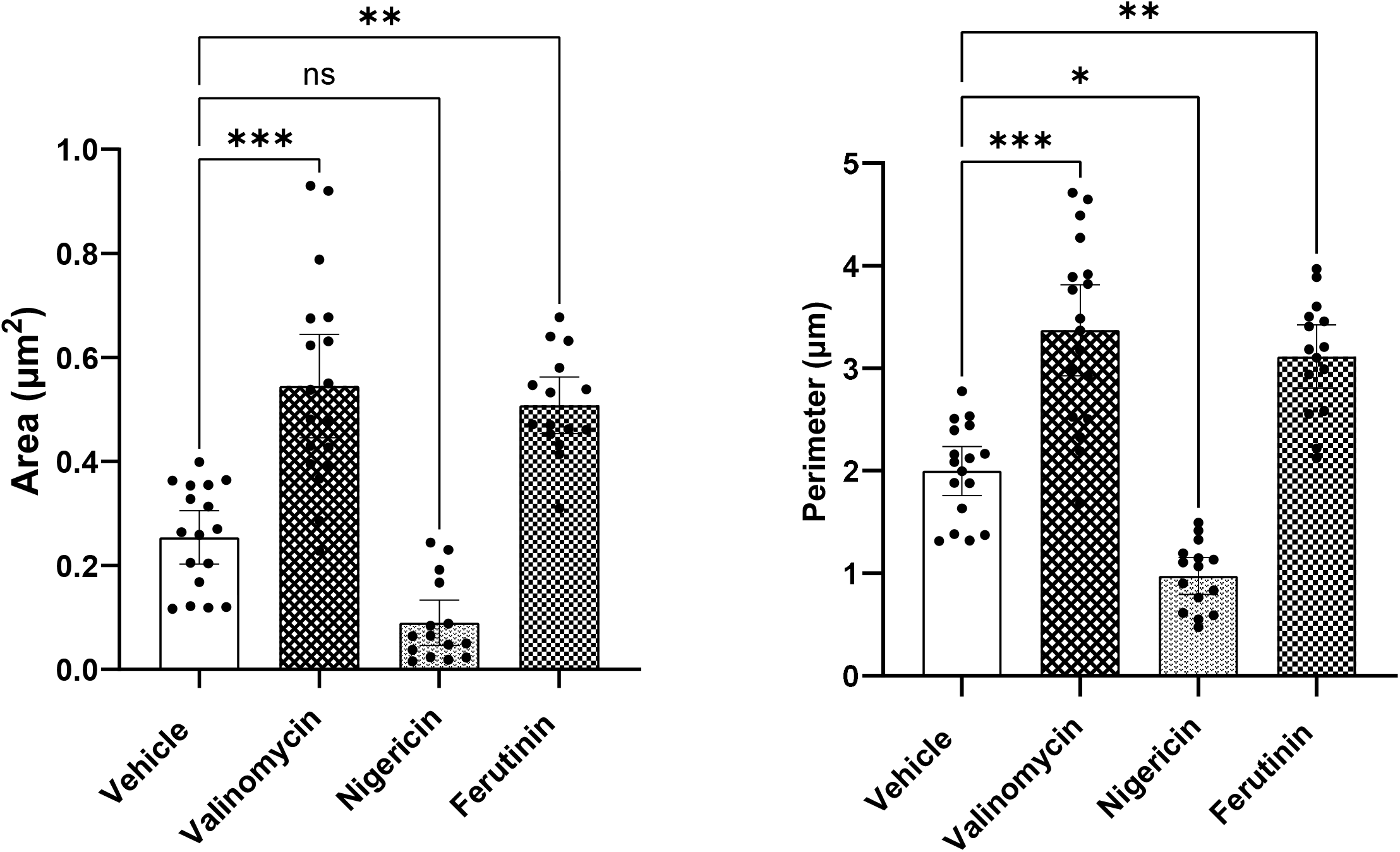
Quantification of the EM images demonstrate increase in mitochondrial area (A) and perimeter (B) following valinomycin and ferutinin treatment indicating organelle swelling.

**Supplementary Figure 2.**
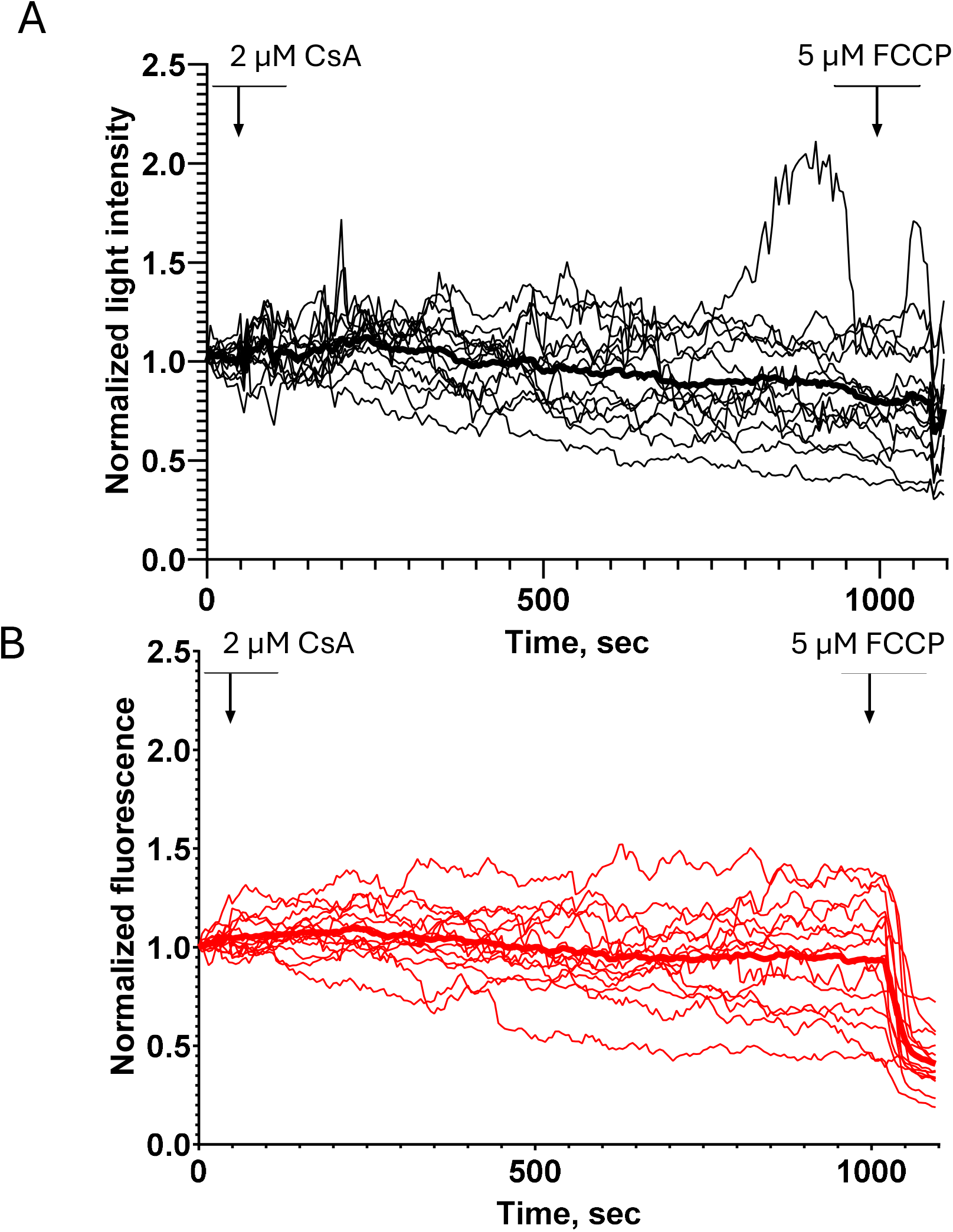
Light scattering (A) and membrane potential (B) measurements of the HAP1 wild-type cells. Note lack of membrane depolarization.

**Supplementary Figure 3.**
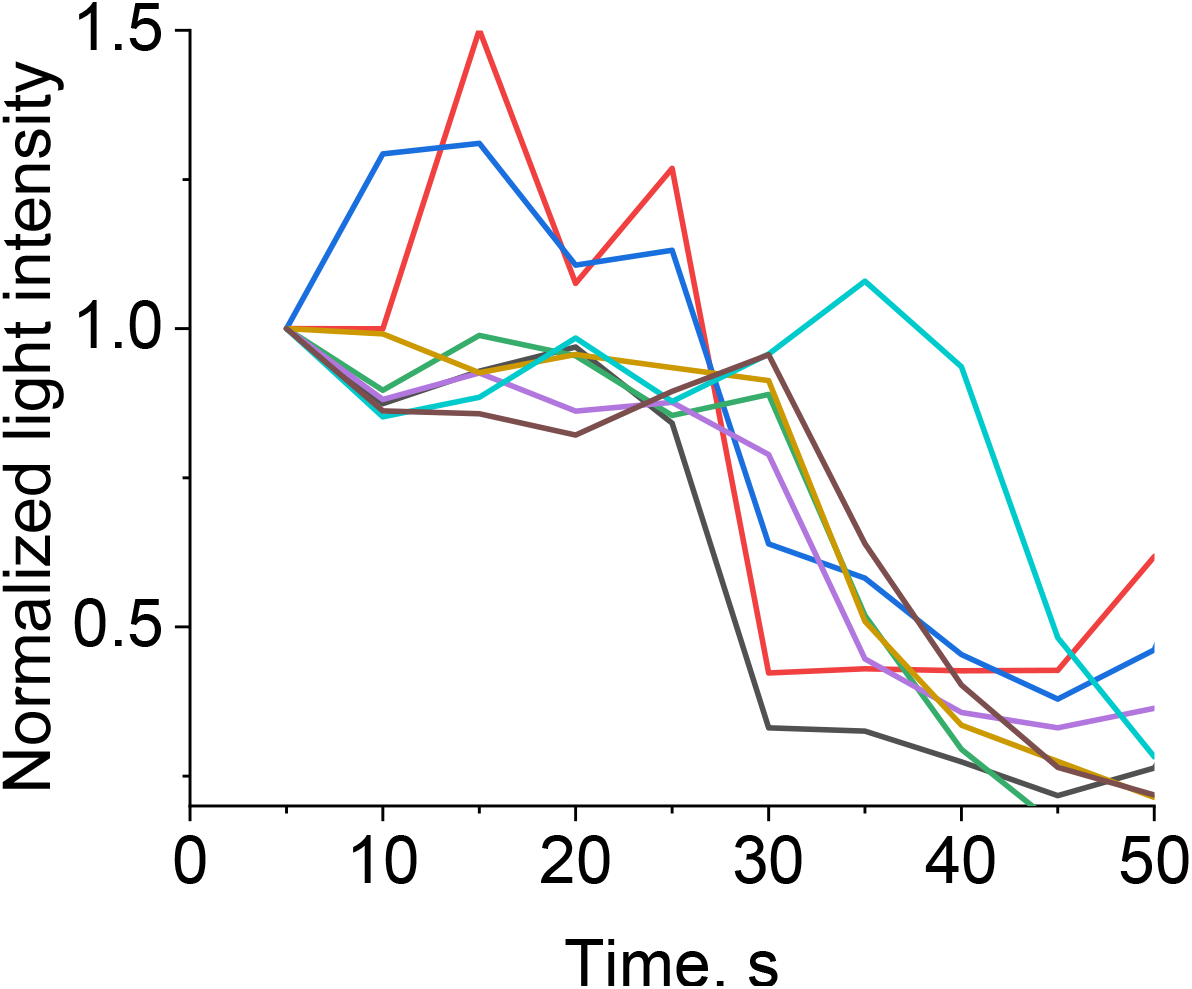
Heterogeneous PTP dynamics in a single cell. Each trace originates from different ROI taken within the same cell. Note the onset of PTP occurs at different timepoints in some populations complete as early as at 30 sec (black trace) and as late as at 50 sec (magenta trace).

## Notes

### Competing Interest Statement

The authors have declared no competing interest.

